# Population estimates of shorebirds on the Atlantic Coast of southern South America generated from large-scale, simultaneous, volunteer-led surveys

**DOI:** 10.1101/2024.02.26.582090

**Authors:** Fernando A. Faria, Joaquín Aldabe, Juliana B. Almeida, Juan J. Bonanno, Leandro Bugoni, Robert Clay, Julian Garcia-Walther, Agustina M. González, Arne Lesterhuis, Guilherme T. Nunes, Nathan R. Senner

## Abstract

Population abundance and trend estimates are crucial to science, management, and conservation. Shorebirds, which are abundant in many coastal habitats and play important roles in coastal ecosystems, are facing some of the most dramatic population declines of any group of birds globally. However, accurate and up-to-date population estimates are lacking for most shorebird species. We thus conducted comprehensive, simultaneous, and community scientist-led surveys of the entire Atlantic Coast of southern South America — stretching from central Brazil to Tierra del Fuego — to gather counts of shorebirds that we combined with remote sensing analyses and two-step hurdle models that accounted for presence and abundance. Our objectives were to estimate shorebird densities by habitat, identify high-concentration areas, understand the environmental factors affecting their distributions, and provide population estimates for both Nearctic and Neotropical species. We counted a total of 37,207 shorebirds of 17 species and, from those counts, estimated that nearly 1.1 million shorebirds use the region’s coastline. We also found that shorebirds occurred in the highest densities in shallow water wetland habitats and that fewer shorebirds occupied areas that were further away from estuaries. Although not directly comparable, our results suggest the population sizes of the Nearctic species whose nonbreeding ranges are predominantly in southern South America may have declined substantially since previous estimates. At the same time, our study represents the first empirically derived population estimates for Neotropical breeding shorebird species and indicates that they are far more abundant than previously thought. Taken together, our results highlight the power of community scientists to carry out structured protocols at continental scales and generate critical data for a group of at-risk species.

## INTRODUCTION

Population size estimates form the basis for understanding many ecological and evolutionary processes (Magurran et al. 2010, Ripple and Breschta 2012). Population size estimates are also vital for management and conservation, as they provide baseline data that can serve as a reference point for evaluating the success of on-the-ground actions (Mace et al. 2008, Monzón and Friedenberg 2018). When applied at large spatial scales, these estimates can help identify species’ distributional patterns and biodiversity hotspots, and even be incorporated into broader analyses of ecosystem function and environmental health (Şekercioğlu et al. 2004, Stuart-Smith et al. 2013). Nonetheless, robust population estimates remain difficult to generate and a lack of such estimates hinders our ability to forecast species’ vulnerabilities to declines and extinctions (Fraser et al. 2022).

Coastal ecosystems support high levels of biodiversity and are among the most productive ecosystems globally (Mitsch and Gosselink 2015). Simultaneously, coastal wetlands are one of the most threatened environments on the planet (Schuerch et al. 2018, IPCC 2022). Habitats such as saltmarshes, estuaries, and coastal lagoons are facing a multitude of threats, including pollution (Naidoo et al. 2015, Hitchcock and Mitrovic 2019), sea level rise (Schuerch et al. 2018, Fagherazzi et al. 2020), and urban development (Lee et al. 2006, Freeman et al. 2019). In fact, most of world’s megacities are located in coastal areas and they continue to attract more people to these regions (Seto 2011, Brown et al. 2013). Increasing urbanization, in turn, constitutes one of the main causes of habitat fragmentation and loss in coastal areas (Liu et al. 2016). Consequently, coastal dependent organisms have been heavily impacted, and monitoring their populations is critical to assessing their conservation status and the health of coastal ecosystems (Studds et al. 2017, Avila et al. 2018, MacNeil et al. 2020).

Among coastal species, birds are thought to be good sentinels of ecosystem health, since they are visible, have broad public appeal, occupy a variety of habitats and trophic levels, and tend to respond quickly to environmental changes (Smits and Fernie 2013). Coastal birds, and especially shorebirds in the order Charadriiformes, are also among the most threatened groups of species in the world (Simmons et al. 2015), with many North American breeding shorebird populations exhibiting accelerating declines that have exceeded 50% over the past few decades (Smith et al. 2023). The existence of these trend estimates, however, belies the fact that we do not have good population estimates for many Nearctic breeding shorebirds and do not know how close they may actually be to extinction (Andres et al. 2012).

The Atlantic coast of southern South America supports the majority of the hemispheric and global populations of a number of Nearctic-breeding migratory shorebird species (Andres et al. 2012). Population estimates from the region were generated for these species nearly four decades ago (Morrison & Ross 1989), but are now in urgent need of updating. Such updates are best developed from data collected in southern South America because: i) many Nearctic species are sparsely distributed during the breeding season across remote parts of the Arctic (McCarty et al. 2020), and ii) rapid turnover rates make estimates harder to generate at migratory stopover sites (Wang et al. 2022). Importantly, many Neotropical species also breed and occupy the coasts of southern South America throughout the year (del Hoyo et al. 1996), but no such empirically derived estimates exist for them (Wetlands International 2023). Generating robust population estimates for both Nearctic and Neotropical species in the region is thus critical to developing targeted conservation efforts.

The limited number of range-wide population estimates that exist for shorebirds, as well as other taxa, can be partially explained by the difficulties posed by coordinating large-scale efforts (i.e., in terms of financial and/or human resources; Stroud et al. 2006). However, recent methodological advances can now facilitate the collection and analysis of data at necessarily large scales. For example, the development of remote sensing technologies has improved ecological studies and conservation actions *via* the enhancement of analytical tools and the quality and resolution of images available across vast geographic regions (Hansen et al. 2013, Nagendra et al. 2013, Rajah et al. 2019). More importantly, the dramatic increase in community science projects that draw participants from the general public has allowed the collection of large datasets that can be used to investigate the distributions (Newson et al. 2016, Schubert et al. 2019), behavior (Robbins et al. 1986) and long-term population trends of birds (Horns et al. 2018, Gillings et al. 2019). Combined, such approaches now make feasible large-scale studies that can generate shorebird population estimates across their nonbreeding ranges. In this context, we conducted the first community scientist-driven, simultaneous, comprehensive survey of shorebirds along the Atlantic Coast of southern South America in 2019. By using a combination of on-the-ground surveys and remote sensing analyses, our objectives were to: 1) create habitat-specific density estimates; 2) identify areas with high concentrations of shorebirds; 3) determine the environmental factors influencing shorebird distributions; and, 4) generate large-scale population estimates for both Nearctic and Neotropical species. Given the rates at which shorebirds are thought to be declining, we hope that these results can be used to guide management actions and decisions while also providing baseline data for future estimates of population trends.

## METHODS

### Study Area and Habitats

Our study area encompassed the southeastern portion of the South American Atlantic Coast — stretching from the Brazilian state of Santa Catarina (28°48′S) along the entire coast of Uruguay and Argentina to the province of Tierra del Fuego (54°65′S) — a total distance of 6,575 km. We included in our surveys any shorebird habitat within 5 km of the coastline. We then divided the study area into nine major regions (two in Brazil, two in Uruguay, and five in Argentina) based on biogeographic boundaries (Spalding *et al*. 2007) and previous survey efforts (e.g., Morrison and Ross 1989; Fig. 1), which allowed some level of interdecadal comparison of our population estimates.

**Fig. 1.**
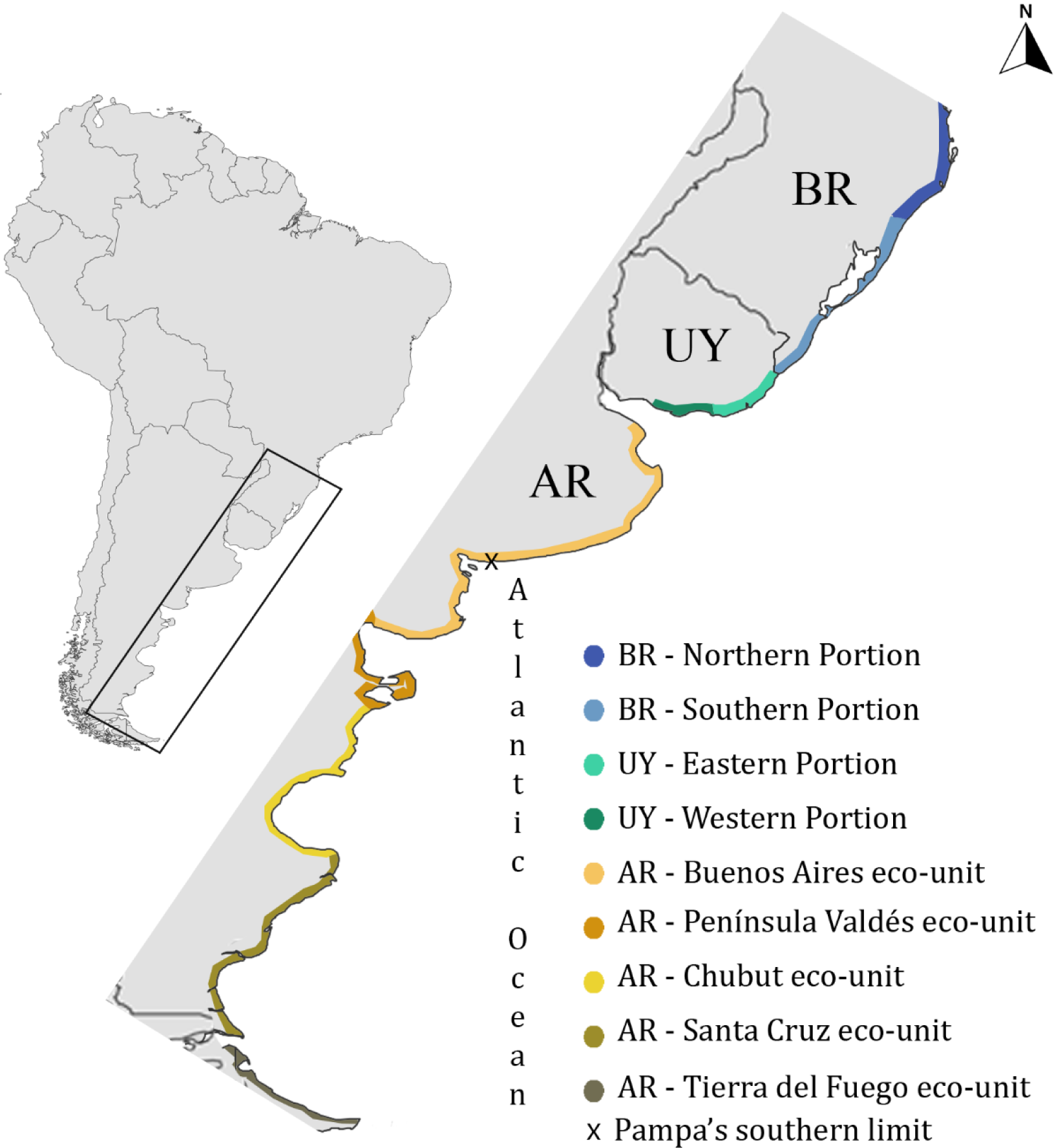
Map of southeastern South America where simultaneous shorebird surveys were conducted from 20 – 29 January, 2019. Distinct colors indicate division of coastal regions used in the sampling design. Colored region width is illustrative and does not represent the 5 km buffer applied to the coast in the analyses. AR – Argentina, UY – Uruguay and BR – Brazil.

As a first step, we chose habitats and sites to survey by relying on previous information indicating their potential importance for migratory and resident shorebirds (Morrison and Ross 1989, Isacch and Martínez 2003, Bencke et al. 2006, Alfaro and Clara 2007, WHSRN 2023), searching the eBird database (Sullivan et al. 2014), and following the methodology developed by Senner and Angulo-Pratolongo (2014) and García-Walther et al. (2017). The habitats included rocky and sandy beaches, *restingas* (broad rocky platforms extending to the lower intertidal zone; Morrison and Ross 1989), intertidal mudflats, shallow water wetlands, and adjacent low vegetation height grasslands. The last two habitat types only occurred in the Pampas Biome, which ranges from the Rio Grande do Sul state in Brazil to the southern tip of the Buenos Aires province in Argentina (Fig. 1).

### Volunteer Training

Before our surveys began, we hosted five workshops — one in Brazil, one in Uruguay, and three in Argentina — to guarantee that all participants received standardized training on shorebird identification, count methods, and data submission through the eBird website. During the workshops, we formed teams of observers based on the number of sites to be surveyed and the degree of experience of the participants, ensuring that all teams included at least one experienced observer. Each team then received georeferenced maps with pre-determined polygons and transects to be surveyed. Surveys were conducted by teams of at least two observers.

### Shorebird Surveys

In order to minimize the potential for birds to move among sites and inflate our estimates, we carried out all surveys within a country during a single 2-d window, while across all countries, we carried out the surveys within a single 9-d window. We were able to survey all known areas with wetland habitats across all three countries. Within each of these areas, we surveyed at least one representative sample of each of the six habitat types — e.g., shallow water wetlands, low vegetation height grasslands, intertidal mudflats, restingas, rocky beaches, and sandy beaches — that were present. Away from these known wetland areas, we randomly chose 244 sandy beach, 15 rocky beach, and 52 *restinga* segments to additionally survey.

Our surveys were conducted following a standardized protocol that accounted for: survey day and hour; area covered (ha) or transect length (m); tidal conditions, when necessary (i.e., within ± 3 h of low tide); and number of observers. We also accounted for habitat-specific detection probabilities by tailoring our survey methods to each habitat type. Thus, following Senner and Angulo-Pratalongo (2014), in grassland and shallow water sites, observers walked the borders of 0.1 × 0.4-km pre-determined polygons using binoculars to count all shorebirds and identify them to species level. For intertidal mudflats, observers walked a pre-defined transect bisecting the habitat and stopped every 0.4 km to perform unlimited time point counts during which they counted and identified every shorebird within a 0.2-km radius. The results from these habitats are presented as birds.ha^-1^. In *restingas* and along rocky and sandy beaches, observers walked pre-determined 0.5-km length transects counting and identifying all birds from the coastline to the habitat’s supralittoral limits. The results from these habitats are presented as birds.km^-1^. Birds crossing transects or polygons in flight were marked separately and – in addition to birds identified to the genus level – not considered in our analyses.

### Spatial Analysis

Because we were unable to survey all of the shorebird habitat within each known wetland area, and because patches of shorebird habitat might exist outside of the previously known wetland areas, we obtained freely available Sentinel-2 satellite images (http://glovis.usgs.gov) to assess the extent of habitats that could be classified as suitable and, potentially, used by shorebirds across the entire study area. Sentinel-2 satellites provide high-resolution (∼10 m^2^) multispectral imagery with 13 bands in the visible, near infrared, and shortwave infrared parts of the light spectrum (Immordino et al. 2019). We opted for cloud-free images from the closest date to the survey period available (29 December 2018 to 22 February 2019). We analyzed each country separately and subset Argentina into two regions because of its large size (Fig. 1). Subsetting helped to increase the overall classification accuracy by reducing the numbers of land-cover types and spectral variation (Bhattarai and Giri 2011).

For the habitats with transect-based surveys, we divided the coastline into 0.5-km length sections. Then, because no good remote sensing tools exist to differentiate among the habitats systematically, we visually classified each section as *restinga*, rocky beach, sandy beach, or ‘unsuitable’ (artificial buildings such as harbors/piers, industries, and cliffs). For the habitats with polygon-based surveys, we generated multispectral images with spectral bands 8, 4 and 3, as near infrared is efficient at differentiating among vegetation types as well as detecting exposed soil/water (Stratoulias et al. 2015, Faria et al. 2021). In addition, we created polygons bordering all estuaries and cities (e.g., areas with >10 ha of buildings and streets rather than simply areas within legal jurisdictional limits) along the coastline. We then used supervised classification models based on a maximum-likelihood algorithm to classify and distinguish among habitat types. This process involved translating the pixel values of a satellite image into distinct habitat categories (Horning et al. 2010) and allowed us to use our actual, on-the-ground, surveyed sites as ‘training sites’ for the model.

We performed a first classification to detect waterbodies inside the 5-km buffer from the coastline. Then, we applied a 1-km buffer around these waterbodies — to restrict low vegetation height grasslands to this buffered area (see Faria et al. 2023) — and classified the remaining habitats to determine their extent. For each polygon/transect classified as suitable, we calculated the i) central coordinates, ii) distance to the nearest city, and iii) distance to the nearest estuary. For polygons, we also calculated the size in hectares of each classified habitat. All GIS analyses were performed in ArcMap 10.8.

### Population Estimates

In order to develop population estimates for the entire study area that accounted for both surveyed and unsurveyed patches of habitat, we had to account for the (potentially) separate factors influencing occupancy (i.e., the presence/absence of a species) and abundance. To do this, we used ‘hurdle’ generalized linear models, which involve two steps: an initial ‘binomial’ step that can be used to evaluate the influence of predictor variables on a species’ presence within a survey area, and a second ‘continuous’ step to evaluate factors influencing a species’ abundance. Models were fitted separately for each country and survey design (i.e., transect or polygon-based surveys) in the R Programming Environment (v. 4.3.2; R Core Team 2020) using the ‘hurdle’ function in the *pscl* package (Jackman 2015). In both portions of the model, we included the predictor variables: species identity, habitat type, distance to nearest city, and distance to nearest estuary. Models for the polygon-based surveys also included habitat size (in ha) as a predictor variable. Before we included them in the models, we standardized all numerical variables using the ‘rescale’ function in the *lme4* package (Bates et al. 2010). Model selection was based on a stepwise procedure using Akaike Information Criterion (AIC; Burnham and Anderson 2002).

Selected models were used to predict the abundance, and 95% confidence intervals around that abundance, of shorebirds in each habitat and country with the ‘prediction’ function in the *marginaleffects* package (Arel-Bundock et al. 2023). Finally, in order to generate site- and species-specific population estimates, we combined results from the ground surveys with the predicted modeled values (Table S1). Surveys that i) did not follow the protocol, or ii) presented an exceptionally high number of individuals were included in final estimates, but not in the models (see below).

## RESULTS

Between 20–29 January 2019, 189 volunteers conducted 387 surveys across the three countries. We counted a total of 37,207 shorebirds from 17 species (11 Nearctic and 6 Neotropical) and 4 families (Charadriidae, Scolopacidae, Haematopodidae, and Recurvirostridae). In total, we estimated from these counts that ∼1.1 million Nearctic and Neotropical shorebirds used the Atlantic Coast of southern South America (Table 1).

**Table 1.**
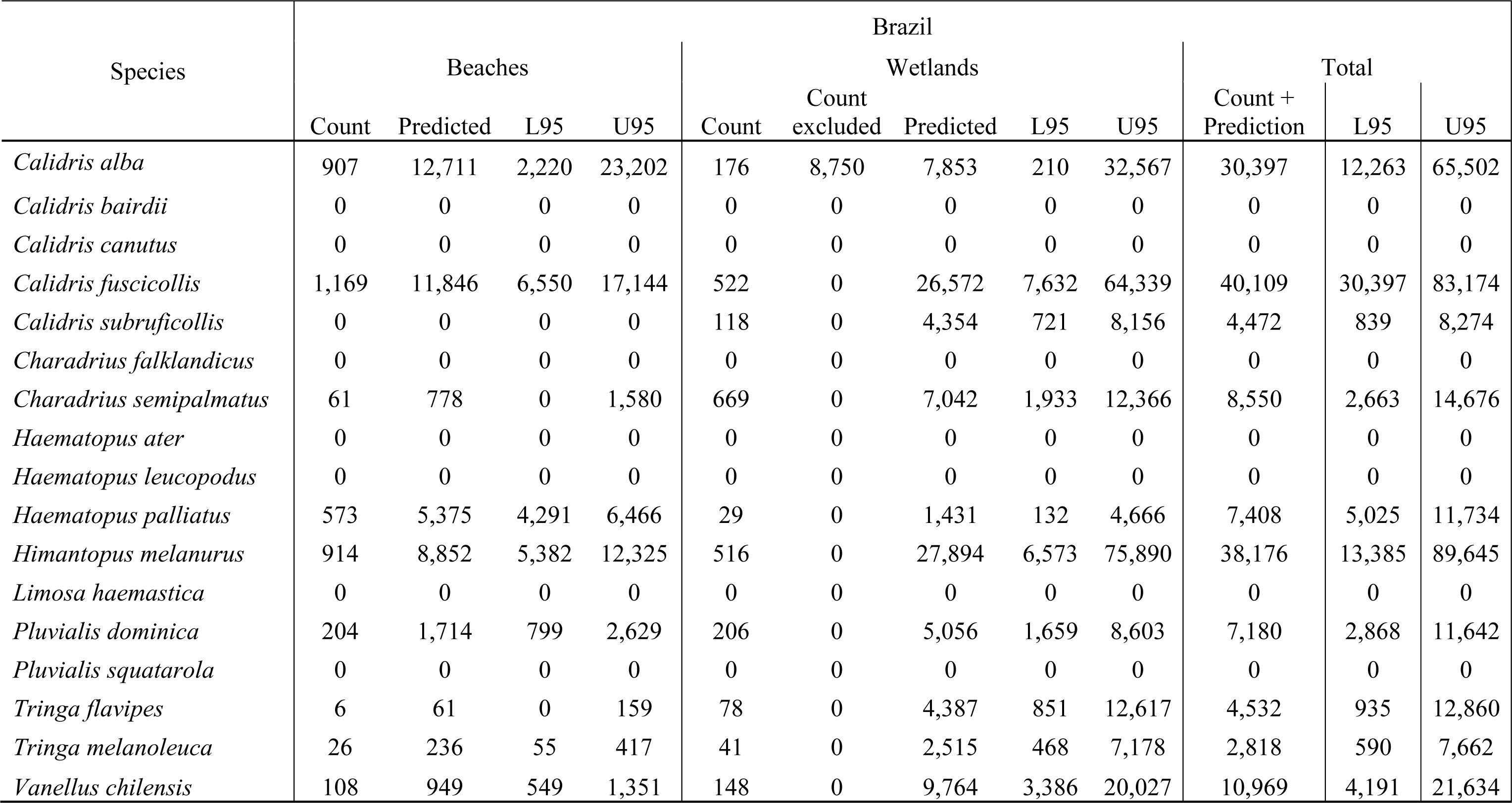
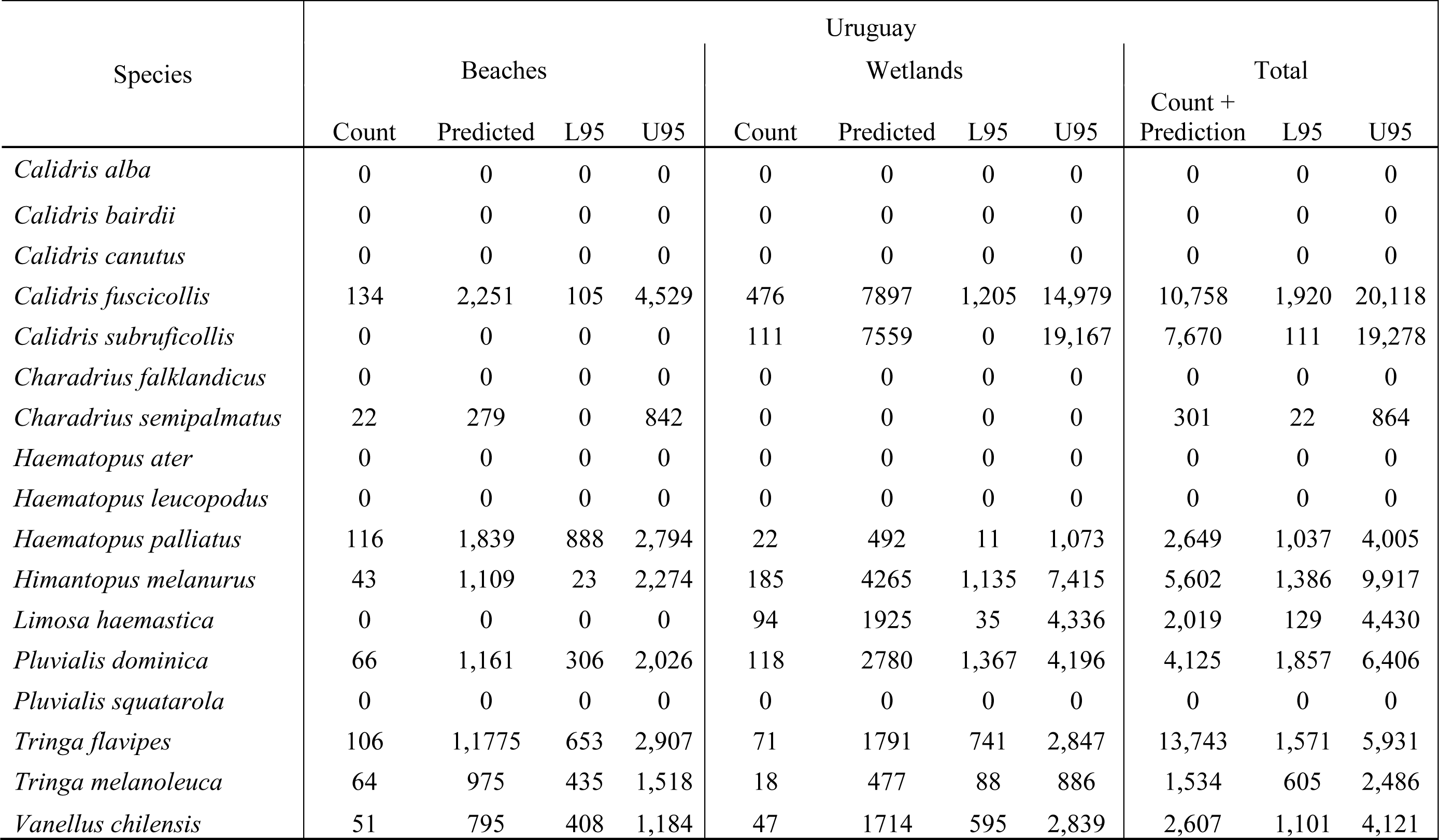
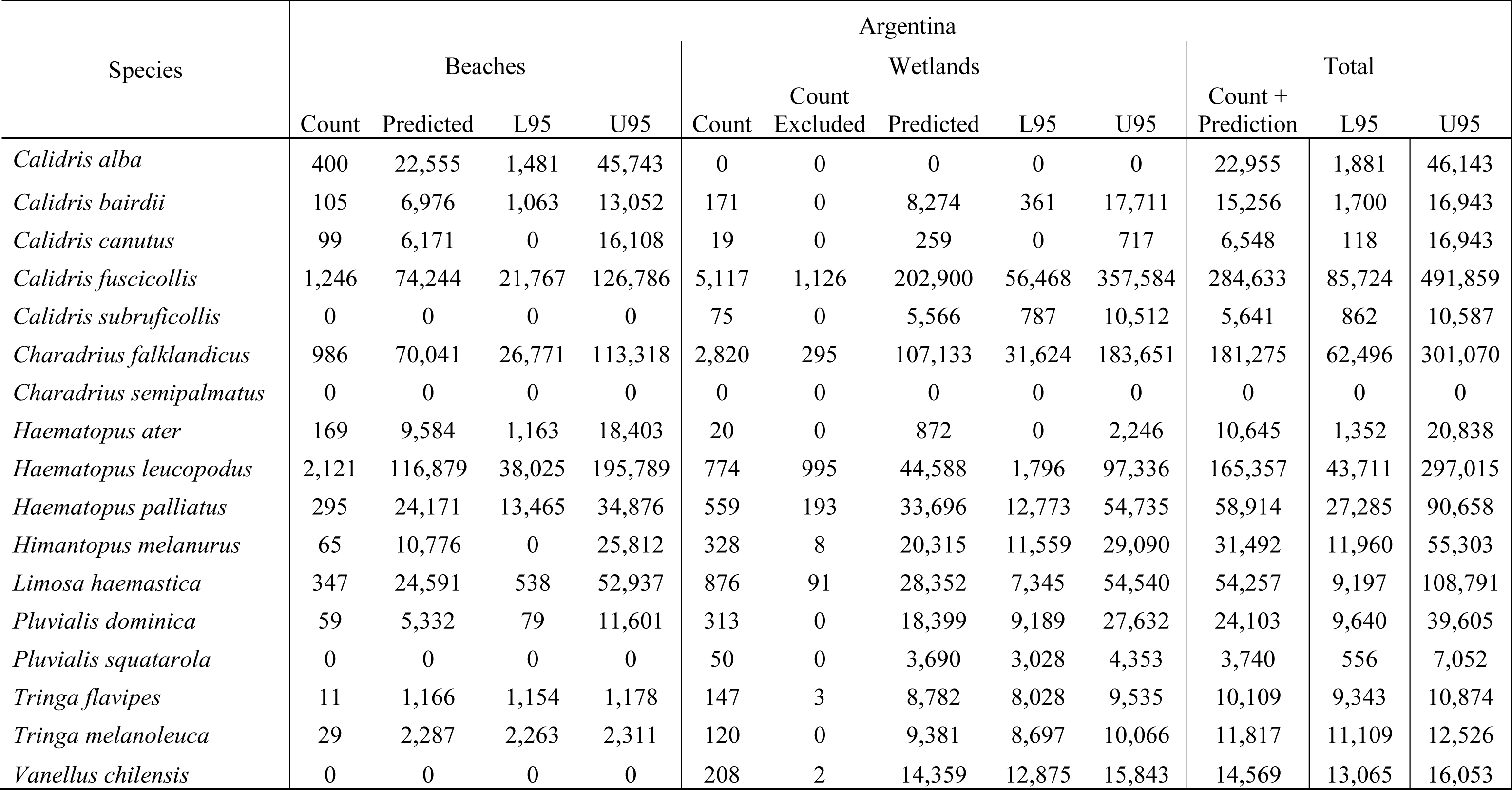
Results of survey and population estimates of Nearctic and Neotropical shorebirds in southeastern South America in January 2019.

Most shorebirds were found in shallow water wetlands, low vegetation height grasslands, and intertidal mudflats (59.8% of individuals). Argentina supported 81.9% of all shorebirds, followed by Brazil (14.0%) and Uruguay (4.1%). Overall, the most abundant species were the Nearctic-breeding White-rumped Sandpiper (*Calidris fuscicollis*), with an estimated 335,500 individuals (95% CI: 118,041 – 595,151), and two Neotropical-breeding species, the Two-banded Plover (*Charadrius falklandicus*) with 181,275 individuals (95% CI: 62,496 – 301,070), and the Magellanic Oystercatcher (*Haematopus leucopodus*) with 165,357 individuals (95% CI: 43,711 – 297,015).

In general, the northern portion of the region was important for sandy beach specialists, such as Sanderling (*C. alba*) and American Oystercatcher (*H. palliatus*), while the southern portion supported higher abundances of intertidal mudflat species such as Hudsonian Godwits (*Limosa haemastica*) and Red Knots (*C. canutus*), as well as species that rely on rocky intertidal habitats, such as Magellanic Oystercatcher (*H. leucopodus*). Consequently, species exhibited distinct distributional patterns, varying from species that were predominantly restricted to the northern portion of the study area — Semipalmated Plover (*C. semipalmatus*), Buff-breasted Sandpiper (*C. subruficollis*), and Black-necked Stilt (*Himantopus melanurus*) — to those restricted to the southern portion of the study area — Magellanic Oystercatcher and Red Knot — and, finally, a few widespread species — American Oystercatcher and White-rumped Sandpiper (Fig. 2).

**Fig. 2.**
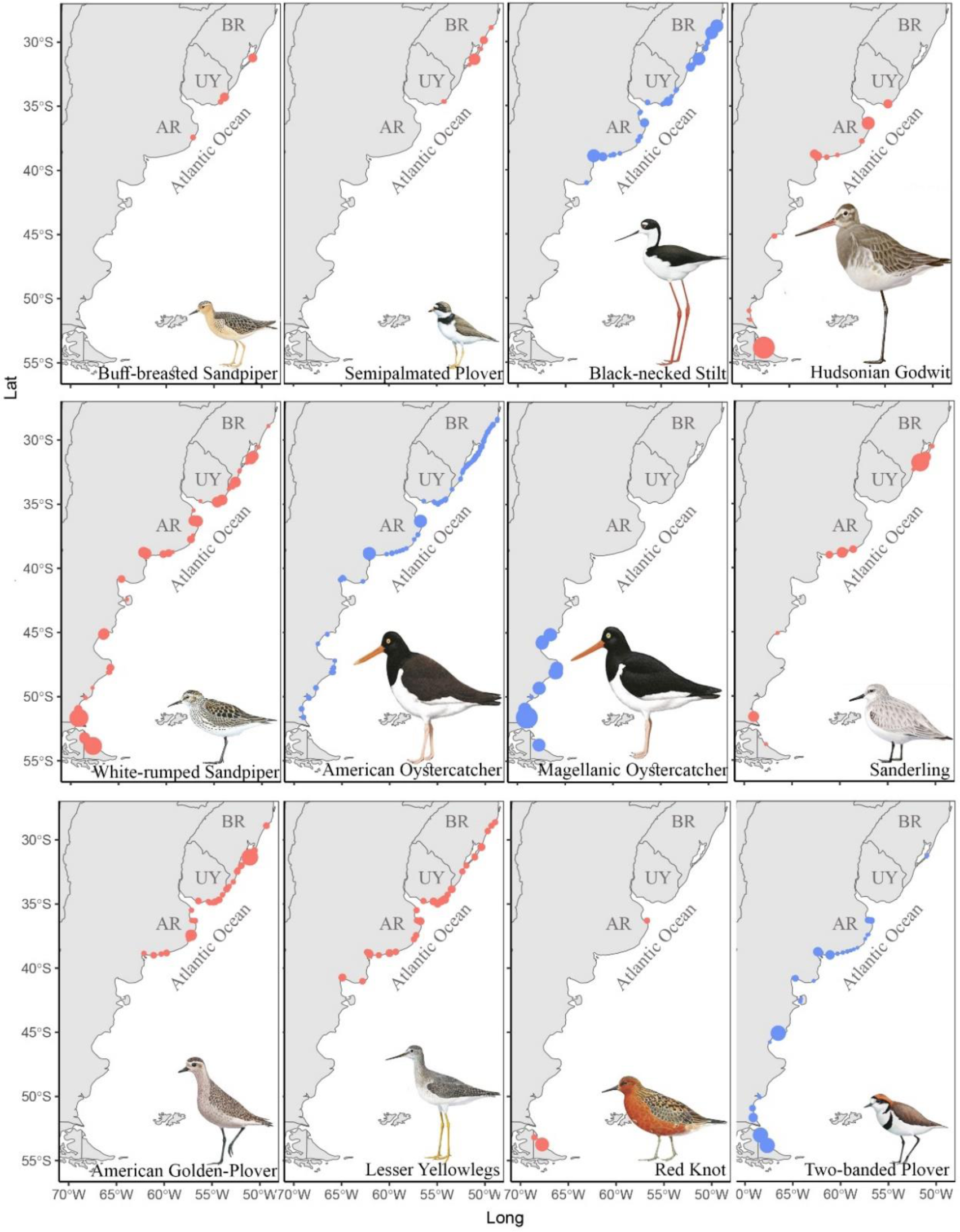
Relative distribution of Neotropical (blue) and Nearctic (red) shorebirds counted during the simultaneous surveys conducted between 20 – 29 January, 2019. The size of points is not directly comparable between maps. AR – Argentina, BR – Brazil and UY – Uruguay.

Our models additionally indicated that increasing distances to estuaries had a negative effect on shorebird abundance, especially for beach habitats. However, we did not find a consistent negative effect of proximity to cities on either shorebird presence or abundance (Fig. 3, Tables 2 & 3). Our species-specific results are presented in Supplemental Document 1.

**Fig. 3.**
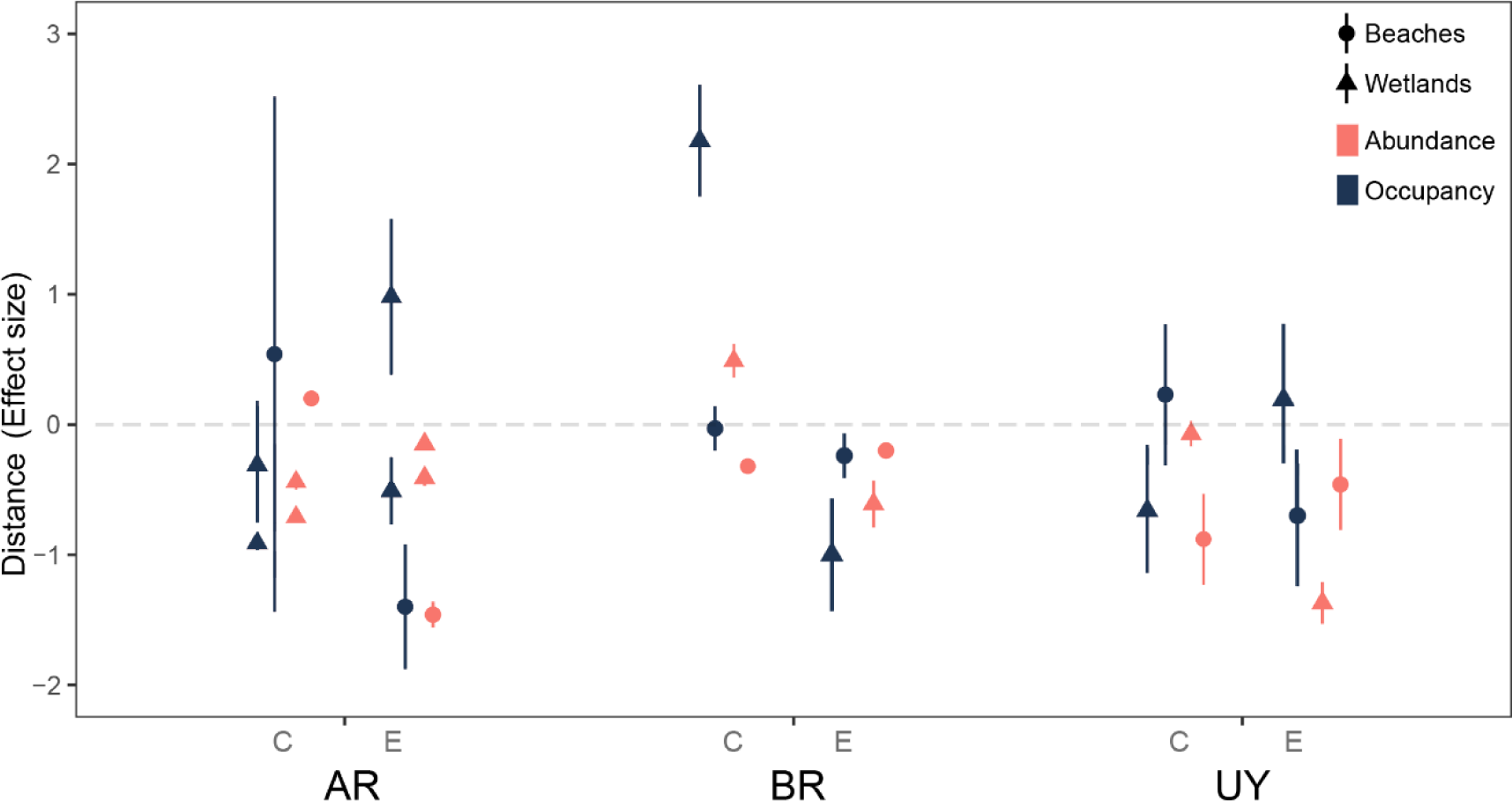
Predicted Effects of distance to closest city (C) and closest estuary (E) on abundance and occupancy of shorebirds counted in beaches and wetlands during simultaneous surveys conducted between 20 and 29 January 2019. The y-axis represents the estimate effect and standard deviation of variables in the three countries: AR – Argentina, BR – Brazil and UY – Uruguay.

**Table 2.** Effects of distance to nearest cities and estuaries on the abundance and occupancy of Nearctic and Neotropical shorebirds using beach habitats in southeastern South America, 2019. SE = standard errors. Bold values represent values of *p* <0.05.

| Beaches | Brazil |  |  | Argentina |  |  | Uruguay |  |  |
| --- | --- | --- | --- | --- | --- | --- | --- | --- | --- |
|  | Estimate | SE | *p* value | Estimate | SE | *p* value | Estimate | SE | *p* value |
| Estuaries |  |  |  |  |  |  |  |  |  |
| Occupancy | -0.25 | 0.17 | 0.15 | -1.41 | 0.48 | **0.003** | -0.71 | 0.54 | 0.18 |
| Abundance | -0.20 | 0.04 | **<0.001** | -1.46 | 0.1 | **<0.001** | -0.46 | 0.35 | 0.18 |
| Cities |  |  |  |  |  |  |  |  |  |
| Occupancy | -0.03 | 0.16 | 0.83 | 0.54 | 1.98 | **0.006** | 0.23 | 0.58 | 0.69 |
| Abundance | -0.32 | 0.04 | **<0.001** | 0.20 | 0.02 | **<0.001** | -0.88 | 0.35 | **0.01** |

**Table 3.** Effects of distance to nearest cities and estuaries on the abundance and occupancy of Nearctic and Neotropical shorebirds using coastal wetland habitats in southeastern South America, 2019. SE = standard errors. Bold values represent values of *p* <0.05.

| Wetlands | Brazil |  |  | Argentina - Region 1 |  |  | Argentina - Regions 2 to 5 |  |  | Uruguay |  |  |
| --- | --- | --- | --- | --- | --- | --- | --- | --- | --- | --- | --- | --- |
| Estuaries | Estimate | SE | *p* value | Estimate | SE | *p* value | Estimate | SE | *p* value | Estimate | SE | *p* value |
| Occupancy | -1.15 | 0.45 | **<0.001** | -0.52 | 0.26 | **0.04** | 0.97 | 0.6 | 0.1 | 0.18 | 0.58 | 0.7 |
| Abundance | -0.61 | 0.18 | **<0.001** | -0.15 | 0.04 | **<0.001** | -0.41 | 0.06 | **<0.001** | -1.37 | 0.16 | **<0.001** |
| Cities | Estimate | SE | *p* value | Estimate | SE | *p* value | Estimate | SE | *p* value | Estimate | SE | *p* value |
| Occupancy | 2.18 | 0.43 | **<0.001** | -0.91 | 0.03 | **0.002** | -0.31 | 0.45 | 0.49 | -0.66 | 0.48 | 0.16 |
| Abundance | 0.49 | 0.13 | **<0.001** | -0.44 | 0.06 | **0.001** | -0.71 | 0.04 | **<0.001** | -0.07 | 0.1 | 0.5 |

## DISCUSSION

Shorebirds are one of the most threatened groups of birds on the planet, yet we lack robust population estimates for most of them (Andres et al. 2012, Smith et al. 2023). Here, we present large-scale population estimates for 17 Nearctic and Neotropical shorebird species from the Atlantic Coast of three countries in southern South America. Our population estimates were mostly lower than previous estimates for Nearctic species (Andres et al. 2012), but higher for Neotropical species (Wetlands International 2012). In addition, we found that both the presence and abundance of shorebirds in beach habitats were higher closer to estuaries. Taken together, these results complement information about recent Nearctic shorebird population declines, while underscoring the substantial knowledge gaps that exist about Neotropical species and highlighting the power of community scientists to carry out complex protocols across large spatial scales.

In general, we found that shorebirds used countries and habitats in similar ways to those detailed in previous studies. Along the Brazilian coast, for instance, Neotropical species were widespread throughout the study area, while Nearctic species occurred mostly along the country’s southern coast, with the highest concentrations of both groups found around Lagoa do Peixe. Furthermore, as with previous studies, the oceanic beaches from Lagoa do Peixe southward to the Uruguayan border were especially important for sandy beach specialists like Sanderlings (Resende and Leewenberg 1987, Vooren and Chiaradria 1990). In Uruguay, we corroborated previous studies, which had shown that Nearctic species were largely restricted to the country’s eastern coast where there are a number of sizeable coastal lagoons (e.g., lagunas Rocha and Merín; Alfaro and Clara 2007, Aldabe et al. 2023). In Argentina, we found the highest shorebird concentrations in previously reported important shorebird areas, such as Bahía Samborombón (Mártinez-Curci et al. 2017), Bahía Blanca (Blanco et al. 2006), the Estuario de Río Gallegos (Ferrari et al. 2002), and Tierra del Fuego (Morrison and Ross 1989, Baker et al. 2005).

Ours are not the first large-scale surveys of Nearctic shorebirds in the region. In 1982, Morrison and Ross (1989) carried out aerial surveys of the exact same coastline. Surprisingly, our estimates were generally higher than those obtained from their surveys. It is possible that this discrepancy is a methodological artifact, because i) their survey methods resulted in lower detection probabilities and/or ii) they categorized most individuals into size and not species-specific groups, making direct comparisons difficult. Even so, in the Brazilian region covered by both surveys, our estimates of small peeps (*Calidris* spp.) were 2.6× higher than their estimates (Morrison and Ross 1989). The same was true for our estimates from Uruguay (3.6× higher for small shorebirds) and, especially, Argentina (7.5× higher for small shorebirds). Some species are notoriously difficult to detect during aerial surveys, especially small grassland shorebirds, whose detectability decreases considerably at distances greater than a few tens of meters (e.g., Buff-breasted Sandpiper; Aldabe et al. 2019, Faria et al. 2023). Other studies using aerial surveys, however, have also found larger numbers of Nearctic shorebirds than Morrison and Ross (1989). For example, 9,710 shorebirds were estimated from Bahía Samborombón, Argentina in 2014, a number 2.9× higher than that found by Morrison and Ross in 1982 (Martínez-Curci et al. 2019).

One species, Red Knots, however, exhibited notable declines in comparison to the counts from Morrison and Ross (1989). This population’s decline has been well documented and resulted in their listing as a threatened species in the United States (USFWS 2021) and an endangered species in Canada (COSEWIC 2007). Despite covering many sites with historically large concentrations — including Bahía San Sebastian and Rio Grande on Tierra del Fuego — our results suggest that, along the Argentine coast, the population has experienced a nearly 75% decline since the early 1980s and now numbers ∼6,500 individuals. These declines have been variously linked with climate change, as well as a decline in food resources and an increase in disturbance and habitat loss at migratory stopover sites, including sites within southeastern South America (González et al. 2006). Continued conservation efforts targeted at this species throughout its range are clearly critical.

In comparison with other, more recent, population estimates of Nearctic breeding shorebird populations, our results for the Nearctic species for which we covered the majority of their wintering ranges (*e.g*., Black-bellied Plovers, and White-rumped and Baird’s *C. bairdii* Sandpipers) unfortunately mirror our results for Red Knots. In general, our estimates were <20% of those proposed by Andres *et al*. (2012) a decade ago. Our results are therefore in line with other studies indicating that most Nearctic migratory shorebird species are currently facing accelerating declines (Smith et al. 2023). That said, for most species, and even for those that spend the nonbreeding season exclusively in southern South America (e.g., Buff-breasted Sandpiper), we may have missed some important sites if they occurred more than 5 km away from the coast (Faria et al. 2023, Aldabe et al. 2023). Increased efforts focused on inland wetlands – especially those along the base of the Andes Mountains in the southern provinces of Chubut and Santa Cruz, and those in the altiplano of the northern provinces of Salta and Jujuy in Argentina – would be helpful in this regard.

In contrast, our estimates for Neotropical breeding species, with the exception of Black-necked Stilt and Blackish Oystercatcher *H. ater*, were higher than previous estimates, even for species whose distributions extend outside of our study area (Wetlands International 2017). For example, our estimates for the coastal-dependent Magellanic Oystercatcher – without considering the Pacific portion of the species’ range – were 1.65× higher than previous estimates. This discrepancy likely stems from the fact that previous estimates were generated almost entirely using the best guesses of experienced researchers, as no large-scale efforts have previously been made to generate reliable data on most Neotropical shorebirds (Wetlands International 2012).

Beyond our population estimates, our results indicate that shorebirds tended to occur closer to estuaries, especially in Brazil and Argentina. Previous studies have shown that beaches and intertidal mudflats adjacent to estuaries support higher densities of potential prey and are key shorebird habitats (Barter 2002, Canham et al. 2021). However, in habitats such as coastal grasslands, close proximity to estuaries can provide additional benefits. For example, grasslands adjacent to estuaries or large waterbodies can receive nutrient inputs and have a more reliable water supply, enhancing soil fertility and potential prey biomass (Lenhart et al. 2015). Some shorebird species are also relative generalists in their habitat use, relying simultaneously on food resources from sandy beaches, mudflats, and grasslands (Faria 2023). Consequently, estuarine complexes with a variety of habitats found in close proximity to each other could reduce the energetic costs of movements between distinct feeding and roosting sites. In this sense, the proximity of non-estuarine habitats to estuaries should be considered when prioritizing shorebird-related management decisions.

Surprisingly, with the exception of wetlands in Brazil, we did not find evidence that the proximity of a given site to urban areas had a significant effect on either shorebird presence or abundance (Tables 2 & 3). We had predicted that a close proximity to cities would have a negative effect on both presence and abundance, especially for shorebirds using beach habitats, as several studies have previously indicated that proximity to humans affects shorebirds during the breeding (Liley and Sutherland 2007, Hevia et al. 2023), migratory (Pfister et al. 1992, Murchison et al. 2016), and nonbreeding seasons (LeDee et al. 2008, Palacios et al. 2022, Swift et al. 2023). However, most cities are located close to estuaries (Small and Nicholls 2003) and, thus, the reliance of shorebirds on estuarine habitats may be stronger than the pressure to avoid urban areas. Other factors may additionally influence shorebird habitat use, such as the distance between roosting and foraging sites (Rogers et al. 2006), predator densities (Goss-Custard et al. 1991, Whitfield 2003), and levels of disturbance from dogs and people that might be related to a city’s size (Maguire et al. 2018). More focused research with a study designed specifically to investigate the influence or urban habitats on shorebirds is thus needed in South America to better answer this question.

It is also noteworthy that our estimates were made possible by the efforts of nearly 200 community scientists. The potential for community scientists to carry out large-scale surveys of shorebird populations is not new to either the region or the Western Hemisphere, as such efforts go back decades (Howe et al. 1989, Lopez-Lanus and Blanco 2004). In our case, though, by relying on community scientists, we were able to both comprehensively cover every known wetland area in the region and survey all of those sites (largely) simultaneously — something that had never previously been attempted on the Atlantic Coast of southern South America, but complements similar efforts implemented on the continent’s Pacific Coast over the preceding decade (Senner and Angulo-Pratolongo 2014, Garcia-Walther et al. 2017). Additionally, our focus on training community scientists allowed us to make use of habitat-specific survey protocols and pre-defined survey areas, generating density estimates that made possible our extrapolations to unsurveyed (and previously unknown) patches of habitat.

Ultimately, our large-scale, volunteer-based study generated updated and much-needed information about shorebirds along the Atlantic Coast of southern South America and should result in the revision of current population estimates for a range of Nearctic and Neotropical species. Our results raise concerns, given the small global shorebird population estimates previously generated by Andres et al. (2012) and the population declines detailed by Smith et al. (2023). Nonetheless, our estimates for some Nearctic species were higher than those found in the region in 1982 (Morrison and Ross 1989), providing some hope that shorebirds can be resilient in face of environmental changes and habitat loss. In combination with our findings about the influence of the proximity of estuaries and urban areas on shorebird distribution, we expect that our results can be used to support species assessments for country-wide red lists, guide management actions, identify areas for conservation action, and serve as a new baseline for future population monitoring and conservation-action evaluation.

## Acknowledgments

The authors are especially grateful to all of the volunteers that conducted the surveys. We also appreciate the statistical help of A. Zeileis and J. Lyons. Core funding was provided by a Neotropical Migratory Bird Conservation Act Grant # F17AP00657 from the U.S. Fish and Wildlife Service to RC and NRS. The following institutions provided additional funding for the project: Aves Argentinas; Aves Uruguay; Área de Proteção Ambiental da Baleia Franca; BirdLife International; CEMAVE/ICMBio; Lagoa do Peixe National Park; Manomet; SAVE Brasil. Funding to FAF was from the *Conselho Nacional de Desenvolvimento Científico e Tecnológico - CNPq*, through the ‘Programa de Pós-Graduação em Oceanografia Biológica’ (FURG). LB is a research fellow at the Brazilian CNPq (Proc. No. 310145/2022-8).

