## Supplementary Material for "Population estimates of shorebirds on the Atlantic Coast of southern South America generated from large-scale, simultaneous, volunteer-led surveys"

Supplemental Document 1

***Neotropical shorebirds***

**American Oystercatcher *Haematopus palliatus***

This species was found in all three countries, occurring mainly in beaches but also in wetlands. The highest predicted proportion of the species population was found in Argentina (85.6%), followed by Brazil (10.7%). Mean densities in beach habitats ranged from 3.3 ind.km$^{-1}$ in Uruguay to 6.6 ind.km$^{-1}$ in Brazil and 7.9 ind.km$^{-1}$ in Argentina. The total population estimate was 68,791 birds (95% CI: 33,347 – 106,397).

**Magellanic Oystercatcher *Haematopus leucopodus***

In contrast to *H. palliates*, this oystercatcher was found only in Argentina and occurred mostly on beaches (38.8 ind.km$^{-1}$), but also wetlands (3.9 ind.ha$^{-1}$). The Magellanic Oystercatcher was the most abundant species among the Haematopodidae in the study area, with an estimated of 165,357 individuals (95% CI: 43,711 – 297,015).

**Blackish Oystercatcher *Haematopus ater***

Similar to *H. leucopodus*, this species was found only in Argentina. However, unlike the other two *Haematopus* species, it occurred predominantly on beaches (91.6%). It was also the least abundant oystercatcher surveyed, with a mean density of 3.2 ind.km$^{-1}$ along Argentinian beaches. Its total population estimate was 10,645 individuals (95% CI: 1,352 – 20,838).

**White-necked Stilt *Himantopus melanurus***

The only Recurvirostridae shorebird included, this species occurred in all three countries. Unlike oystercatchers, the species was most abundant (~70% individuals) in wetlands in comparison to beaches. Its total population estimate was 75,270 birds (95% CI: 26,731 – 154,865).

**Southern Lapwing *Vanellus chilensis***

This species occurred in all habitats in all regions, except Argentine beaches, with the highest density found in Brazilian wetlands (1.6 ind.km$^{-1}$). [1]. The total population estimate was 28,145 lapwings (95% CI: 18,377 –41,808).

**Two-banded Plover *Charadrius falklandicus***

This species was included only in analyses of Argentine habitats, as just two individuals were found in Brazil. Argentine densities ranged from 9.3 ind.ha$^{-1}$ in wetlands to 23.1 ind.km$^{-1}$ in beaches. The total population estimate was 181,275 individuals (95% CI: 62,496 – 301,070).

***Nearctic shorebirds***

**American Golden-Plover *Pluvialis dominica***

This species was found in all three countries, both along beaches and in wetlands. The highest proportion of individuals was estimated to occur in Argentina (68%), but the highest densities occurred along the Uruguayan and Brazilian beaches (2.1 ind.km$^{-1}$). The total population estimate was 35,408 birds (95% CI: 14,365 – 57,653).

**Black-bellied Plover *Pluvialis squatarola***

Contrary to *P. dominica*, this species occurred mainly in Argentina, with only 10 individuals observed in Brazil. Densities in Argentinian wetlands were estimated at 0.3 ind.ha$^{-1}$ and the total population estimate was 3,740 birds (95% CI: 556 – 7,052).

**Semipalmated Plover *Charadrius semipalmatus***

This plover was restricted to the northern portion of the study area, absent from surveys in Argentina, and with 92.5% of its estimated abundance occurring in Brazil. The total population estimate was 8,851 individuals (95% CI: 2,685 – 15,540).

**Greater Yellowlegs *Tringa melanoleuca***

This species occurred in all countries, both in beach and wetlands. The highest densities were found along Uruguayan beaches (1.8 ind.km$^{-1}$), although the highest proportion of individuals was estimated from Argentina (73.1%). The total population estimate was 16,169 birds (95% CI: 12,304 – 22,674).

**Lesser Yellowlegs *Tringa flavipes***

This species had similar occurrence patterns to Greater Yellowlegs and was present in both beaches and wetlands. As with *T. melanoleuca*, Uruguay was also the country with high density of individuals (20,3 ind.km$^{-1}$ in beach habitats), and it was estimated that

48.4% of individuals occurred in this country. The total population estimate was 28,384 individuals (95% CI: 11,849 – 29,665).

**Buff-breasted Sandpiper *Calidris subruficollis***

This species occurred only in wetlands. Uruguay hosted the highest proportion of individuals (43.2%), followed by Argentina (31.7%) and Brazil (25.1%). Densities were also higher in Uruguay (1.8 $ind.ha^{-1}$), followed by Brazil (0.7 $ind.ha^{-1}$) and Argentina (0.5 $ind.ha^{-1}$). The population estimate was 17,783 individuals (95% CI: 1,812 – 38,139).

**Baird's Sandpiper *Calidris bairdii***

This species was only detected in Argentina, with relatively similar proportions estimated from interior (54.4%) and beach habitats (45.5%). The population estimate was 15,526 individuals (95% CI: 1,700 – 31,039).

**White-rumped Sandpiper *Calidris fuscicollis***

This was the most abundant shorebird in surveys and was present in all three countries, both in beach and wetlands. This also had the highest densities of any species in both beach (4.1 $ind.km^{-1}$ and 24.6 $ind.km^{-1}$) and interior habitats (2 $ind.ha^{-1}$ and 17.6$ind.ha^{-1}$) in Uruguay and Argentina, respectively. In Brazil, densities were 14.3 $ind.km^{-1}$ in beaches and 4.4 $ind.ha^{-1}$in interior habitats. The population estimate was 335,500 individuals (95% CI: 118,041 – 595,151).

**Hudsonian Godwit *Limosa haemastica***

Godwits occurred mainly in Argentina (96.4%), but were also observed in wetlands in Uruguay. The species' density in these habitats was estimated at 2.5 $ind.ha^{-1}$ in Argentina and 0.5 $ind.ha^{-1}$ in Uruguay. The highest density occurred along the Argentine beaches (8.13 $ind.km^{-1}$). The total population estimate was 56,276 individuals (95% CI: 9,326 – 113,221).

**Sanderling *Calidris alba***

This species was the most abundant shorebird in Brazil, mainly in beach habitats, where it was the densest species (15.0 $ind.km^{-1}$). Absent from Uruguay, Sanderlings were also present along Argentine beaches, which hosted 43% of the estimated abundance of the species in the southeastern South America. The total population estimate was 53,352 individuals (95% CI: 14,144 – 111,645).

**Red Knot *Calidris canutus***

This species was only detected during surveys in Argentina. However, contrary to *C. bairdii*, this species was mainly (95.7%) found in beach habitats, with a density of 2.0 ind.km$^{-1}$. The population estimate was 6,548 individuals (95% CI: 118 – 16,943).

**Table S1.** Country and habitat-specific measurements of survey efforts and model predictions of shorebird simultaneous surveys conducted between 20 and 29 January 2019.

| | **Country** | **Habitats** | **Survey Effort** | **Prediction** |
|---|---|---|---|---|
| Beaches (km) | BR | Sandy | 77.5 | 674 |
| | | Rocky | 1.5 | 25.5 |
| | UY | Sandy | 19 | 387 |
| | | Rocky | 6 | 149 |
| | AR | Sandy | 25.5 | 1,889.50 |
| | | Restinga | 26 | 1002 |
| Wetlands (ha) | BR | Grasslands | 25.66 | 13,538.50 |
| | | Mudflats | 27.3 | 540.2 |
| | | Shallow water | 3.19 | 1,329.50 |
| | UY | Grasslands | 44.3 | 3,577.70 |
| | | Shallow water | 60.3 | 507.9 |
| | AR | Grasslands | 80.8 | 15,733.20 |
| | | Mudflats | 931.8 | 118,901.30 |
| | | Shallow water | 100.6 | 9,744.40 |
